## Supplementary Information file for "Single AAV-mediated mutation replacement genome editing in limited number of photoreceptors mediate marked visual restoration"

**Supplementary Information (Nishiguchi KM et al. Single AAV mutation replacement)**

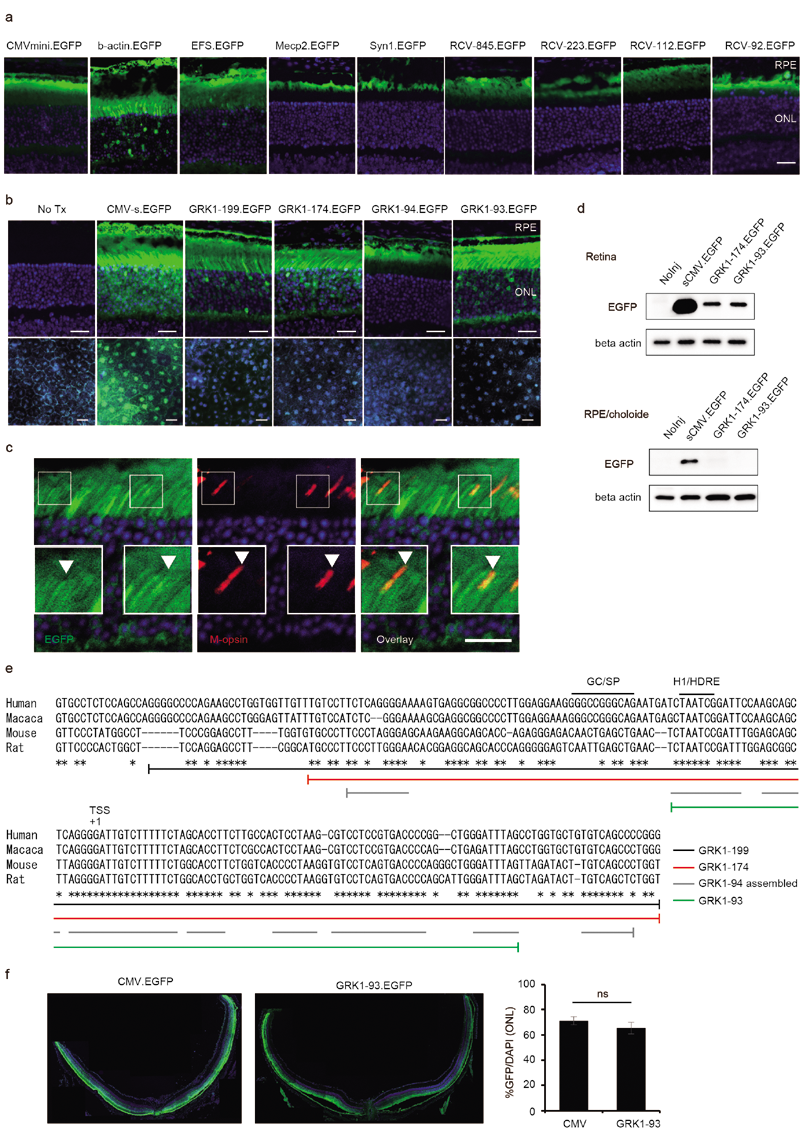

**Fig. S1| Selection of minimal neural retina-specific promoters**

a. *In vivo* EGFP reporter analysis of retinal sections, performed three weeks after AAV injection with various published and modified promoters (N = 2 each). The ONL coincides with the photoreceptor cell bodies. The details of the promoters used are shown in Table S1. The sections were stained only with DAPI.
b. *In vivo* EGFP reporter analysis of *GRK1* promotor deletion mutants in retinal sections (upper panels) and retinal pigment epithelium flat mounts (lower panels). The sections and flat mounts were stained only with DAPI.
c. Immunohistochemistry of cone photoreceptors. *GRK1-93*-driven EGFP and cone-specific M-opsin were co-localized.
d. Western blot analysis of reporter EGFP. Note that *GRK1-93*-driven EGFP was not detected in the RPE.
e. Sequence of the *GRK1* promoter mutants tested in the experiment.
f. Comparison of transduction efficacy, based on reporter EGFP expression (green) driven by a CMV promoter or *GRK1-93* promoter, in histological sections of the eye. The proportion of photoreceptors transduced with AAV (EGFP-positive cells/DAPI positive cells in ONL, N = 3 each) is shown.

ONL, outer nuclear layer; EGFP, enhanced green fluorescent protein; NoInj, not injected. GC/SP, GC-rich regions presumably interacting with Sp protein; H1/HDRE, potential homeodomain (Crx) binding site.

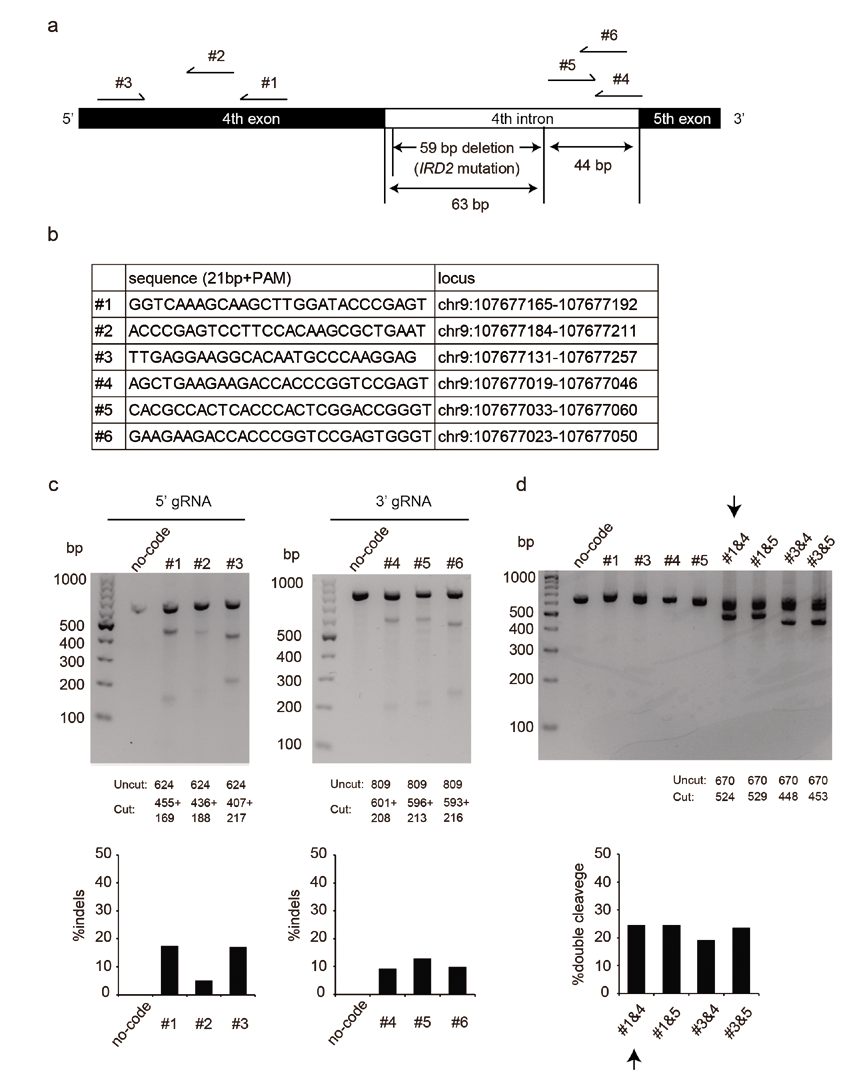

**Fig. S2| Selection of gRNA pair**

a. A schematic map of gRNA designed.
b. List of gRNAs and their sequences.
c. T7E1 assay for each gRNA. The expected DNA size is displayed under the representative gel images from 3 independent replicates, all showing similar results. Quantified editing efficiency is displayed in the lower panels.
d. Electrophoresis of DNA fragments after cleavage of the genome with pairs of gRNAs. Representative images from 3 independent replicates, all showing similar results. Quantified double cleavage efficiency is displayed in the lower panel.

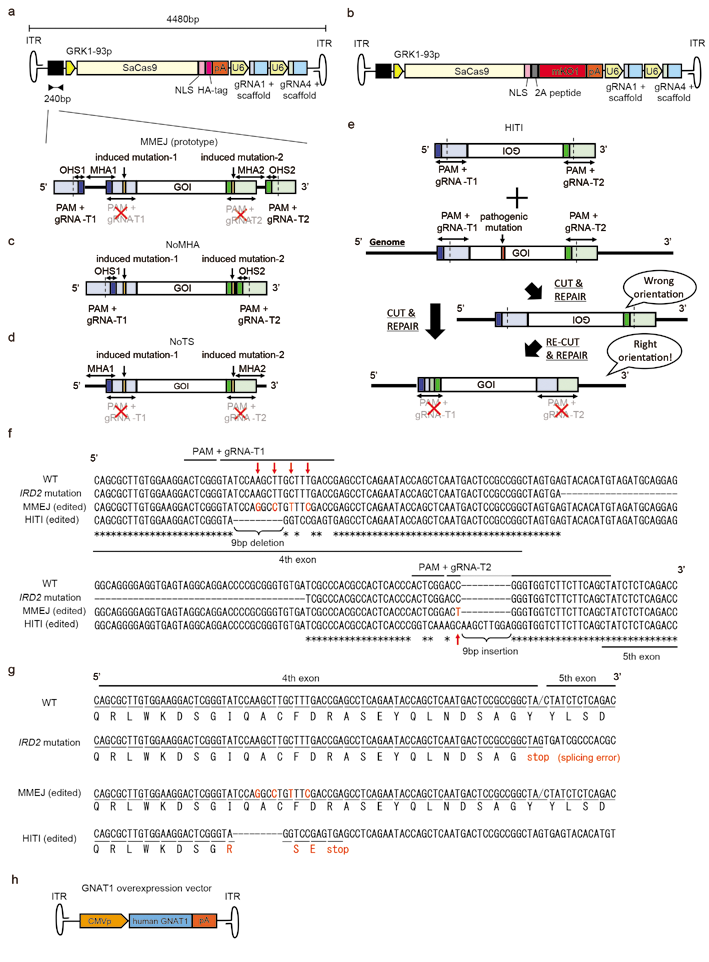

**Fig. S3| Design of genome editing vectors**

a. Design the prototype MMEJ mutation replacement vector (MMEJ vector) with an enlarged map of the donor template. Total size was 4,480 bp.
b. An enlarged map of the MMEJ vector with a reporter for lineage tracing experiments. SaCas9 and mKO1 were linked with 2A peptide. Total size was 5,201 bp.
c. An enlarged map of the donor template without flanking microhomology arms (NoMHA).
d. An enlarged map of the donor template without flanking gRNA target sites (NoTS).
e. An enlarged map of the donor template for HITI mutation replacement vector and an illustration of HITI-mediated mutation replacement strategy. In this approach, GOI was inserted in the opposite direction relative to the flanking gRNA target sites in the donor template, so that when the GOI was inserted into the genome in a correct orientation through NHEJ, these sites would be disrupted, preventing re-cleavage by SaCas9. Conversely, when the GOI were inserted in the genome in a wrong orientation, the flanking gRNA target sites would remain intact, which will be subjected to re-cleavage by SaCas9 until GOI is positioned in the correct orientation.
f. Comparison of editing outcomes at the genome level after successful applications of MMEJ- and HITI-mediated mutation replacement. Induced mutation in the gRNA target sites were highlighted in red and with arrows and nucleotides conserved across 4 sequences displayed was marked with * at the bottom of the sequence alignment. Note, 9bp deletion and 9bp insertion took place at both ends of GOI in HITI.
g. Comparison of genome editing outcomes at amino acid level after successful application of MMEJ- and HITI-mediated mutation replacement. Note, altered amino acids in reference to the wildtype sequence were highlighted in red. As a result of the significant nucleotide alterations of the 5’ gRNA target site after HITI-mediated mutation replacement, 3 missense changes and 9 bp deletion followed by a nonsense mutation took place in the 4^th^ exon.

MMEJ, micro-homology-mediated end-joining; NoMHA, no microhomology arms, NoTS, no target sites; HITI, homology-independent targeted integration; GOI, gene of interest; gRNA-T, guide RNA target; PAM, protospacer adjacent motif; ITR, inverted terminal repeat; MHA, micro homology arm; NLS, nuclear localizing signal; pA, ploy A; PAM, protospacer adjacent motif; mKO1, monomeric Kusabira-Orange 1; WT, wild-type.

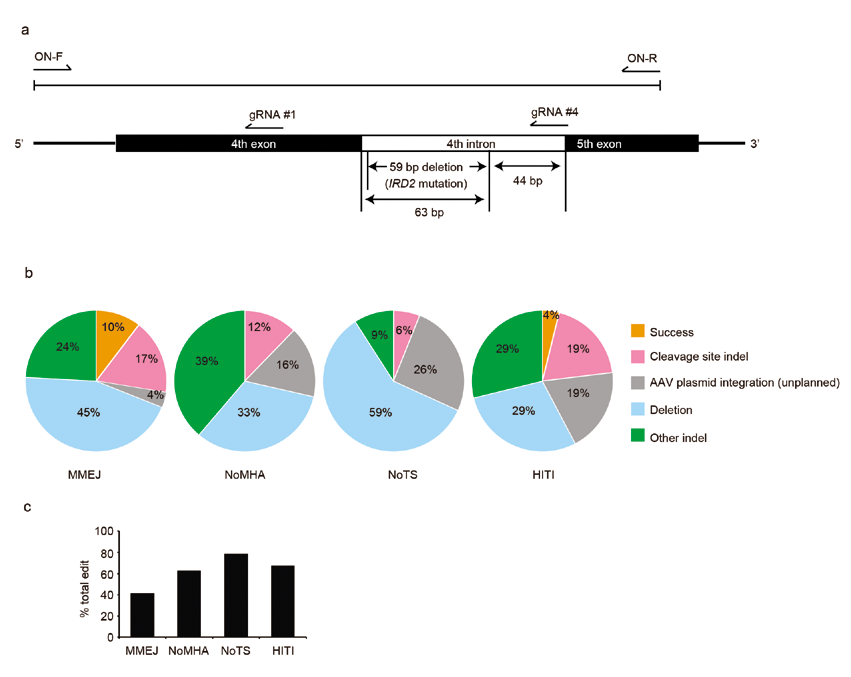

**Fig. S4| *In vitro assessment of on-target site following mutation replacement therapy in cultured murine neural cells***

a. Schematic map of primers used for ON-target site analysis. ON-F and ON-R indicates the position of forward and reverse primer designed on the mouse genome.
b. Breakup of on-target sequencing results of the genome edited clones amplified from murine Neuro2A cell lines after transfection of the mutation replacement vector. Total clones sequenced in this experiment were 70, 67, 84 and 77 for MMEJ, NoMHA, NoTS, and HITI, respectively. The design of each vector is outlined in Fig. S3. Note, “success” indicates successful mutation replacement without induction of unplanned mutations elsewhere.
c. The rate of genome edited clones among sequenced clones.

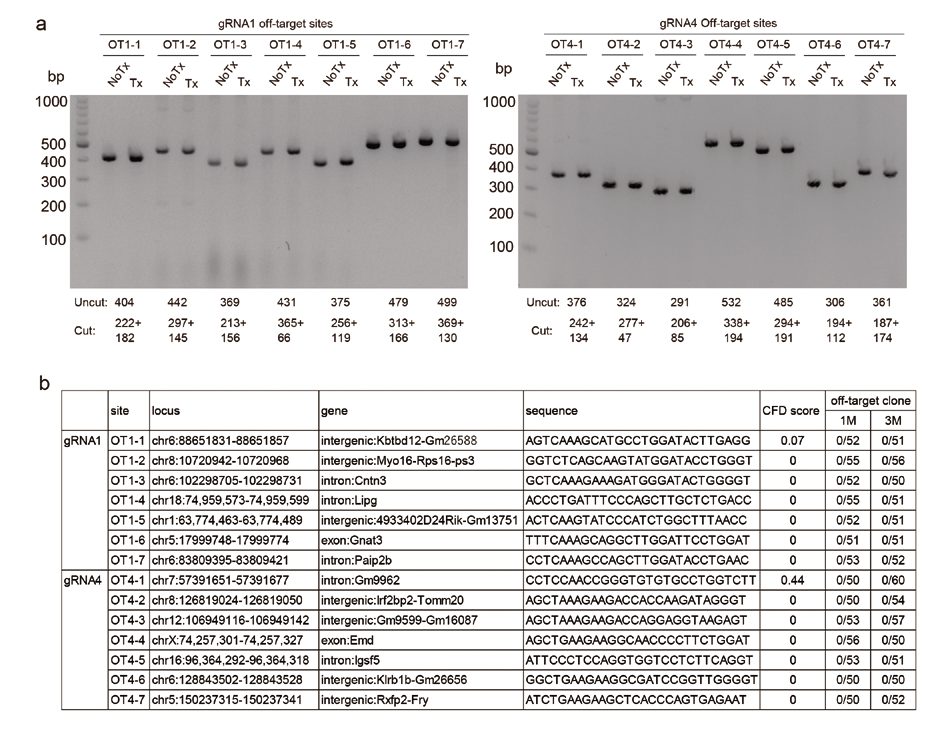

**Fig. S5| Off-target analysis**

a. T7E1 assay of 7 sites for each gRNA (total of 14 sites) predicted with CRISPOR (<http://crispor.tefor.net/>). Expected DNA size before (Uncut) and after (Cut) T7E1 digestion is displayed under representative gel images from 4 independent replicates. Note that there were no bands of the expected sizes in the presence of off-target mutations.
b. Summary of off-target sites and results of Sanger sequencing. “CFD scores” represent the likelihood of off-target DNA damage induction. The numbers of sequenced clones and mutations found (all *“zero”*) are expressed as denominators and numerators, respectively, in the column “off-target clone”. OT, off-target. Tx, treated with T7E1; NoTx, Not treated.

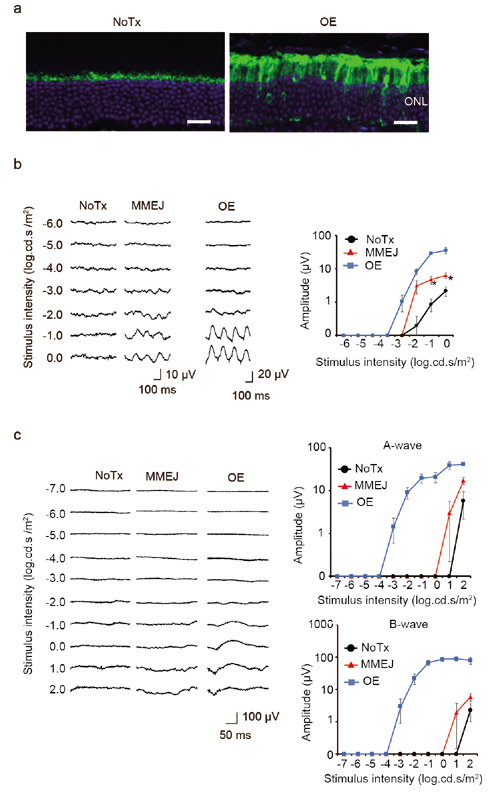

**Fig. S6| Comparison of MMEJ-mediated mutation replacement and gene supplementation therapy in *Pde6c^cpfl1/cpfl1^Gnat1^IRD2/IRD2^* mice**

a. Representative GNAT1 immunohistochemistry images of the retina in *Pde6c^cpfl1/cpfl1^Gnat1^IRD2/IRD2^* mice treated and untreated with *GNAT1* over-expression.
b. 6Hz flicker ERGs recorded from the eyes treated with either MMEJ-mediated *Gnat1-IRD2* mutation replacement (MMEJ, N = 9) or *GNAT1* over-expression (OE, N = 9) and the untreated (NoTx, N = 6) eyes of *Pde6c^cpfl1/cpfl1^Gnat1^IRD2/IRD2^* mice.
c. Single flash ERGs recorded from the eyes treated with either MMEJ-mediated *Gnat1-IRD2* mutation replacement (MMEJ, N = 7) or *GNAT1* over-expression (OE, N = 7) and the untreated (NoTx, N = 6) eyes of *Pde6c^cpfl1/cpfl1^Gnat1^IRD2/IRD2^* mice. Data represent the mean ± S.E.M.; *P < 0.05 for comparison between NoTX and MMEJ. MMEJ, micro-homology-mediated end-joining.

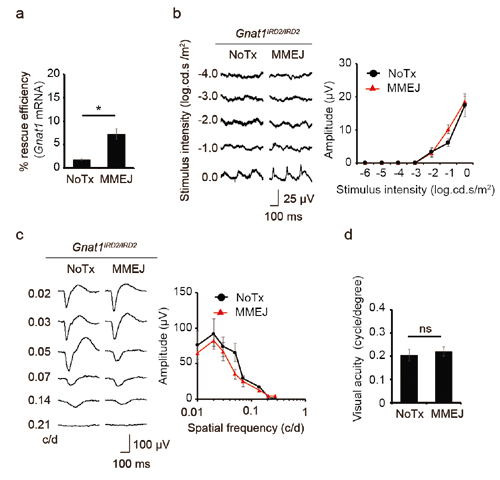

**Fig. S7| Treatment of *Gnat1^IRD2/IRD2^* mice with MMEJ-mediated mutation replacement AAV vector**

**a,** Genome editing efficiency as measured with RT-PCR (N = 5) in *Gnat1^IRD2/IRD2^* mice. **b, c, d**. Flicker ERGs (**b**, N = 7), OKR (**c**, N = 8) and pVEPs (**d**, N = 7) detected no treatment effect. Data represent the mean ± S.E.M.; *P < 0.05; MMEJ, micro-homology-mediated end-joining; NoTx untreated; ns, not significant.
