## Supplementary material for "Single AAV-mediated mutation replacement genome editing in limited number of photoreceptors mediate marked visual restoration": Table S1

| Promoter | Sequence (5'>3') |
|---|---|
| b-actin | GTCGAGGTGAGCCCCACGTTCTGCTTCACTCTCCCCATCTCCCCCCCCTCCCCACCCCCAATTT<br>TGTATTTATTTATTTTTTAATTATTTTATGCAGCGATGGGGGCGGGGGGGGGGGGGGGCGCGCGC<br>CAGGCGGGGCGGGGCGGGGCGAGGGGCGGGGCGGGGCGAGGCGGAGAGGTGCGGCGGCA<br>GCCAATCAGAGCGGCGCGCTCCGAAAGTTTCCTTTTATGGCGAGGCGGCGGCGGCGGCGGCC<br>CTATAAAAAGCGAAGCGCGCGGCGGGCGGGAGTCGCTGCGTTGCCTTCGCCCCGTGCCCCGC<br>TCCGCGCCGCCTCGCGCCGCCCGCCCCGGCTCTGACTGACCGCGTTACTCCCACAG |
| CMV mini | GGTAGGCGTGTACGGTGGGAGGCCTATATAAGCAGAGCTCGTTTAGTGAACCGTCAGATCGCCT<br>GGAG |
| CMV-s | CGTTACATAACTTACGGTAAATGGCCCGCCTGGCTGACCGCCCAACGACCCCCGCCCATTGACG<br>TCAATAATGACGTATGTTCCCATAGTAACGCCAATAGGGACTTTCCATTGACGTCAATGGGTGGA<br>GTATTTACGGTAAACTGCCCACTTGGCAGTACATCAAGTGTATCATATGCCAAGTACGCCCCCTA<br>TTGACGTCAATGACGGTAAATGGCCCGCCTGGCATTATGCCCAGTACATGACCTTATGGGACTTT<br>CCTACTTGGCAGTACATCTACGTATTAGTCATCGCTATTACCatgAGAGGGTATATAATGGAAGCT<br>CGACTTCCAG |
| EFS | TGGCTCCGGTGCCCGTCAGTGGGCAGAGCGCACATCGCCCACAGTCCCCGAGAAGTTGGGGG<br>GAGGGGTCGGCAATTGAACCGGTGCCTAGAGAAGGTGGCGCGGGGTAAACTGGGAAAGTGAT<br>GTCGTGTACTGGCTCCGCCTTTTTCCCGAGGGTGGGGGAGAACCGTATATAAGTGCAGTAGTCG<br>CCGTGAACGTTCTTTTTCGCAACGGGTTTGCCGCCAGAACACAGGT |
| GRK-199 | GGGCCCCAGAAGCCTGGTGGTTGTTTGTCCTTCTCAGGGGAAAAGTGAGGCGGCCCCTTGGAG<br>GAAGGGGCCGGGCAGAATGATCTAATCGGATTCCAAGCAGCTCAGGGGATTGTCTTTTTCTAGC<br>ACCTTCTTGCCACTCCTAAGCGTCCTCCGTGACCCCGGCTGGGATTTAGCCTGGTGCTGTGTCA<br>GCCCCGGG |
| GRK-174 | TGTCCTTCTCAGGGGAAAAGTGAGGCGGCCCCTTGGAGGAAGGGGCCGGGCAGAATGATCTAA<br>TCGGATTCCAAGCAGCTCAGGGGATTGTCTTTTTCTAGCACCTTCTTGCCACTCCTAAGCGTCCT<br>CCGTGACCCCGGCTGGGATTTAGCCTGGTGCTGTGTCAGCCCCGGG |
| GRK-94 | TCTCAGGGGATCTAATCGGATTAGCAGCTAGGGGATTGTCTTTTTCTGCACCTTCTCCTAAGGTC<br>CTCCGTGACCCCGGATTTAGTGTCAGCCC |
| GRK-93 | TCTAATCGGATTCCAAGCAGCTCAGGGGATTGTCTTTTTCTAGCACCTTCTTGCCACTCCTAAGC<br>GTCCTCCGTGACCCCGGCTGGGATTTAG |
| Mecp2 | AGCTGAATGGGGTCCGCCTCTTTTCCCTGCCTAAACAGACAGGAACTCCTGCCAATTGAGGGCG<br>TCACCGCTAAGGCTCCGCCCCAGCCTGGGCTCCACAACCAATGAAGGGTAATCTCGACAAAGA<br>GCAAGGGGTGGGGCGCGGGCGCGCAGGTGCAGCAGCACACAGGCTGGTCGGGAGGGCGGG<br>GCGCGACGTCTGCCGTGCGGGGTCCCGGCATCGGTTGCGCGC |
| Syn1 | CTGTGAGGGGGTTATTTCTCTACTTTCGTGTCTCTGAGTGTGCTTCCAGTGCCCCCCTCCCCCCA<br>AAAAATGCCTTCTGAGTTGAATATCAACACTACAAACCGAGTATCTGCAGAGGGCCCTGCGTATG<br>AGTGCAAGTGGGTTTTAGGACCAGGATGAGGCGGGGTGGGGGTGCCTACCTGACGACCGACC<br>CCGACCCACTGGACAAGCACCCAACCCCCATTCCCCAAATTGCGCATCCCCTATCAGAGAGGG<br>GGAGGGGAAACAGGATGCGGCGAGGCGCGTGCGCACTGCCAGCTTCAGCACCGCGGACAGTG<br>CCTTCGCCCCCGCCTGGCGGCGCGCGCCACCGCCGCCTCAGCACTGAAGGCGCGCTGACGTC<br>ACTCGCCGGTCCCCCGCAAACTCCCCTTCCCGGCCACCTTGGTCGCGTCCGCGCCGCCGCCG<br>GCCCAGCCGGACCGCACCACGCGAGGCGCGAGATAGGGGGGCACGGGCGCGACCATCTGCG<br>CTGCGGCGCCGGCGACTCAGCGCTGCCTCAGTCTGCGGTGGGCAGCGGAGGAGTCGTGTCGT<br>GCCTGAGAGCGCAG |
| RCV-845 | TCAGACATATTGACTCACATCAGCCTCACAATGACAGTGTGGTAGATGCTATGATGCCCATTTATT<br>CAAGAAAGACTTGCTGGGGGCCCAAGCCTATCAGGTTCTGGCAATGGAAATGCATCCGGGAACA<br>ACACAGACAAAATCCCATCCTTCACGGAGCTTTCATTCCGGTGAGGGGACATGCAGCGTGCCGA<br>TGATGGTGGGGGTTGTGCTATTATAGATAGAGGGTCAGTATAGGCCTGGCTGAGCAGGTGACAT<br>TTAAGCAGAGACATGACTGCAGTGAAGGTTTAGAAGAGTGTCCCAGGTGGAAAGAGGTGAAGAA<br>ACAAGCCTGGGGCCACACAGCTAGTTAAGTAGCCAAGCTGGGACTTGAACCCATGCTGGCCCC<br>AGAGTCTGTGCTCCTAACCATTGCATTCTAGGGCTTGATATGAGATGCCAGCCCCGCCCCGAGA<br>TGCTCAGAGTTAGTGAGGAAGAGAAACAGGGAACATGGCTGCTGTTAGAGGGCTGGGGCTGGG<br>GGTGCCGGAGGCCCCAGCTCTGAGGGTTCCAACCTCCTGTCCTGTTCTAGTATCGTCCCGGGA<br>GGCCGAGATGAATTGCCTGCCTGCCCTGGGCTCTTTATTTTAATCTCACTAGGGTTCTGGGAGC<br>ACCCCCCCCCACCGCTCCCGCCCTCCACAAAGCTCCTGGGCCCCTCCTCCCTTCAAGGATTGC<br>GAAGAACTGGTCGCAAATCCTCCTAAGCCACCAGCATCTCGGTCTTCAGCTCACACCAGCCTTG<br>AGCCCAGCCTGCGGCCAGGGGACCACGCACGTCCCACCCACCCAGCGACTCCCCAGCCGCTG<br>CCCACTCTTCCTCACTC |

| | |
|---|---|
| RCV-223 | ACTAGGGTTCTGGGAGCACCCCCCCCACCGCTCCCGCCCTCCACAAAGCTCCTGGGCCCCTC<br>CTCCCTTCAAGGATTGCGAAGAACTGGTCGCAAATCCTCCTAAGCCACCAGCATCTCGGTCTTC<br>AGCTCACACCAGCCTTGAGCCCAGCCTGCGGCCAGGGGACCACGCACGTCCCACCCACCCAG<br>CGACTCCCCAGCCGCTGCCCACTCTTCCTCACTC |
| RCV-111 | ATATTGACTCACATGGCAATGGAAATGCTCTGAGGGTTCCAACCTCCTGTCCTGTTCTGCCTGCC<br>CTGGGCTCTTTATTTTAATCTCACTAGGGCCCCTCCTCCCTTCAAG |
| RCV-92 | ATATTGACTCACAGCTCTGAGGGTTCCAACCTCCTGTCCTGTTCTGCCTGCCCTGGGCTCTTTAT<br>TTTAATCTCACCCCTCCTCCCTTCAAG |
