## Supplementary material for "Single AAV-mediated mutation replacement genome editing in limited number of photoreceptors mediate marked visual restoration": Table S2

| Primer ID | Sequence (5’>3’) | Use |
|---|---|---|
| ON-F (in vitro and in vivo) | CAGATGAAGTGAGTGTTCCCTG | On-target |
| ON-R1 (in vivo) | ACCTTCACTCAAGACACTGACG | On-target |
| ON-R2 (in vitro, single cleavage) | GAGATAGCTGAAGAAGACCACCC | On-target |
| ON-R3 (in vitro, double cleavege) | GCTCAGTGGGCACATATCCTG | On-target |
| OT1-1F | TGCATCTGCCACCTCTGAAG | Off-target |
| OT1-1R | TCACAGTCACAGAACCAGGC | Off-target |
| OT1-2F | TCTCAAGGAGCCCAGAGTGA | Off-target |
| OT1-2R | ACTTACTCGAGGGGCAGCTA | Off-target |
| OT1-3F | ATGCTGAGAGCTGATGCTCC | Off-target |
| OT1-3R | GGTTAACCACAGGGCTCAGG | Off-target |
| OT4-1F | ACACAGCTAAGCATCAGGCAGAG | Off-target |
| OT4-1R | TGTGTCTACCACAAGTGCCC | Off-target |
| OT4-2F | GGGTTTTGAAATACTAACCACATGG | Off-target |
| OT4-2R | AGGTAAGTCTCTTGGGGGAAG | Off-target |
| OT4-3F | TGGTAGCAGATAAGGAAGGGT | Off-target |
| OT4-3R | CAGATGTCCCACAGCTGCTT | Off-target |
